# Population-scale study of eRNA transcription reveals bipartite functional enhancer architecture

**DOI:** 10.1101/426908

**Authors:** Katla Kristjánsdóttir, Yeonui Kwak, Nathaniel D. Tippens, John T. Lis, Hyun Min Kang, Hojoong Kwak

## Abstract

Enhancer RNAs (eRNA) are non-coding RNAs transcribed bidirectionally from active regulatory sequences. Their expression levels correlate with the activating potentials of the enhancers, but due to their instability, eRNAs have proven difficult to quantify in large scale. To overcome this, we use capped-nascent-RNA sequencing to efficiently capture the bidirectional initiation of eRNAs. We apply this in large scale to the human lymphoblastoid cell lines from the Yoruban population, and detected nearly 75,000 eRNA transcription sites with high sensitivity and specificity. We identify genetic variants significantly associated with overall eRNA initiation levels, as well as the transcription directionality between the two divergent eRNA pairs, namely the transcription initiation and directional initiation quantitative trait loci (tiQTLs and diQTLs) respectively. High-resolution analyses of these two types of eRNA QTLs reveal distinct positions of enrichment not only at the central transcription factor (TF) binding regions but also at the flanking eRNA initiation regions, both of which are equivalently associated with mRNA expression QTLs. These two regions - the central TF binding footprint and the eRNA initiation cores - define the bipartite architecture and the function of enhancers, and may provide further insights into interpreting the significance of non-coding regulatory variants.

## Introduction

Regulation of transcription is achieved mainly through binding of transcription factors (TFs) at transcription regulatory elements (TREs), such as promoters and enhancers. Genes are expressed from promoters, which integrate regulatory signals from proximal and distal enhancers to determine the amount of RNA product. Such regulatory networks are key to most cellular processes, including development, cell-type differentiation, and stress response, and their misregulation can cause disease. In fact, a large majority of disease associated genetic variation is estimated to affect TREs (Maurano et al. 2012; Gusev et al. 2014). Therefore, considerable efforts have gone into connecting genetic variation to molecular phenotypes at TREs, and to understand how those might affect gene expression (Kasowski et al. 2010; Degner et al. 2012; McVicker et al. 2013; Banovich et al. 2014; Battle et al. 2015; Garieri et al. 2017; Schor et al. 2017; Ferreira et al. 2016).

Enhancer transcription arises in addition to the target promoter activation in species as diverse as flies and humans (Kim et al. 2010; Kaikkonen et al. 2013; Hah et al. 2013; Andersson et al. 2014; Henriques et al. 2018) and its levels track with enhancer activity (Core et al. 2014; Henriques et al. 2018). A pair of enhancer RNAs (eRNAs) are generally transcribed in opposite directions from core transcription-initiation regions flanking the central transcription-factor-binding site (TFBS) of the enhancer (**Fig. 1A**) (Core et al. 2014; Andersson et al. 2015). While the production of eRNAs has been used to identify active enhancers across numerous cell types and tissues (Andersson et al. 2014), the roles eRNAs play in gene regulation have not yet been elucidated (Lam et al. 2014). Complicating eRNA detection and quantification is the fact that they are rapidly degraded (Andersson et al. 2014); this makes it less suitable to use methods that rely on steady-state RNA, such as RNA-seq or Cap Analysis of Gene Expression (CAGE).

Nascent RNA sequencing methods, such as <u>P</u>recision nuclear <u>R</u>un-<u>O</u>n sequencing with 5′-<u>capp</u>ed (m7G) RNA enrichment (PRO-cap, **Fig. 1A**) (Kwak et al. 2013; Kruesi et al. 2013; Core et al. 2014), overcome this challenge by capturing capped RNA at the synthesis stage.

The mapping of genetic variation to molecular phenotypes at different stages of gene expression has provided important insights into the DNA sequences underlying gene regulation (Pickrell et al. 2010; Majewski and Pastinen 2011; Degner et al. 2012; McVicker et al. 2013; Banovich et al. 2014; Battle et al. 2015; Li et al. 2016; Cannavò et al. 2017). Similarly, mapping genotypes to transcription phenotypes at enhancers will help connect changes at enhancers to changes in gene expression, revealing potential mechanisms for gene regulation. Recent studies have mapped genetic variation to transcription at promoters and enhancers using CAGE, revealing quantitative trait loci (QTLs) associated with alternative promoter usage, promoter shape, and expression (Garieri et al. 2017; Schor et al. 2017). But, the drawbacks of CAGE in quantifying unstable eRNA expression limited comprehensive profiling of enhancer-associated QTLs compared to promoter-associated QTLs. Given the properties of PRO-cap that allow it to detect and quantify unstable transcription, we anticipate that a much more comprehensive list of enhancer-associated QTLs can be identified.

This study leverages the variation in transcription initiation at transcribed TREs (tTREs), measured by PRO-cap in lymphoblastoid cell lines (LCLs) from 67 individuals, to study enhancer architecture and activity. We find thousands of genetic variants that affect either transcription initiation levels (tiQTLs) or the directionality of initiation (diQTLs) at enhancers. We find that these two types of QTLs are enriched at distinct positions within the enhancer architecture. Importantly, both variant types show significant association with mRNA expression, illustrating their potential functionality in gene regulation. Overall, through our genetic analysis investigating the pattern of enhancer transcription, our data reveal a bipartite architecture of enhancers.

## Results

### Capped-nascent-RNA sequencing reveals transcribed regulatory elements with high resolution and sensitivity

**Figure 1.**
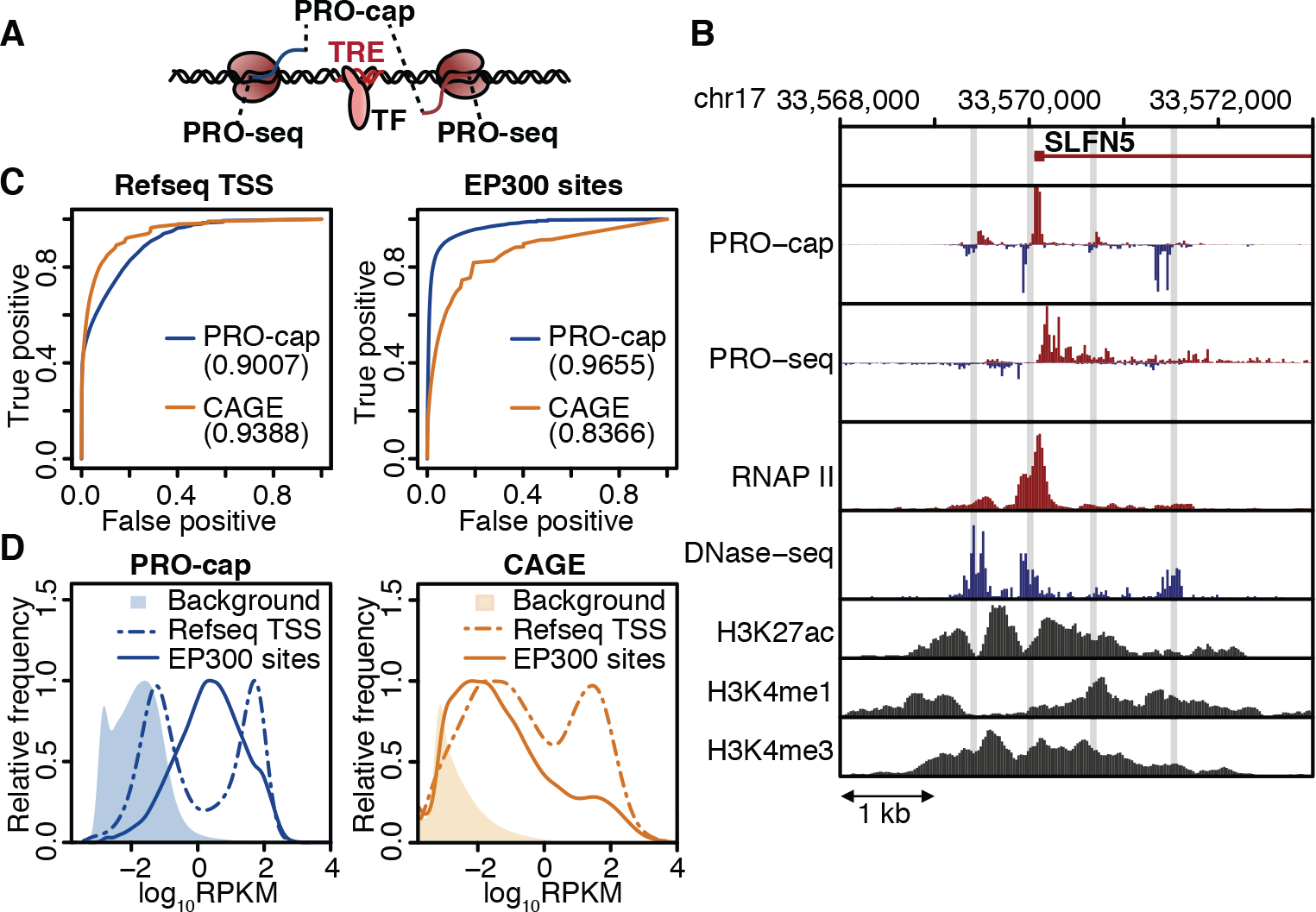
PRO-cap identifies tTREs with high resolution and sensitivity. (**A**) Schematic of bidirectional transcription at tTREs. PRO-cap measures nascent-capped-RNA levels and identifies the precise TSS positions (5′ end); PRO-seq measures the 3′ end of RNAs associated with transcriptionally-engaged polymerase. **(B)** Transcription and chromatin marks at the *SLFN5* locus. PRO-cap, PRO-seq, and DNase-seq data are derived from the YRI LCLs, RNAP II, H3K27, H3K4me2 and H3K4me3 ChIP-seq data are from ENCODE’s LCL, GM12878. Shaded regions indicate PRO-cap-identified tTREs. (**C**) Receiver operating characteristic (ROC) plots of PRO-cap and CAGE at annotated gene promoters (n=12,272) or EP300 bound enhancers (n=18,956). Area under ROC (AUROC) is in parenthesis. (**D**) Histograms of PRO-cap and CAGE log read count distributions at annotated gene promoters and EP300-bound enhancers with background distributions shaded (n=1,000,000).

We prepared PRO-cap libraries from 69 Yoruban lymphoblastoid cell lines (LCLs) (**Supplementary Table 1**) for which a large number of transcriptome and chromatin datasets are available (Pickrell et al. 2010; Degner et al. 2012; McVicker et al. 2013; Banovich et al. 2014; Battle et al. 2015; Li et al. 2016). We combined all PRO-cap datasets and identified transcribed transcriptional regulatory elements (tTREs), including both enhancers and promoters, with bidirectional divergent nascent transcription within 300 basepairs (bp) of one another (**Supplementary Fig. 1A**). We identified 87,826 tTREs (**Supplementary Table 2**) with high sensitivity and resolution, as illustrated by examples at two loci (**Fig. 1B, Supplementary Fig. 1B**). We determined how well our approach identifies Refseq annotated promoters and EP300 bound enhancers and found that it performs substantially better at identifying enhancers than CAGE (ENCODE Project Consortium 2012) (**Fig. 1C, D**).

We separated the tTREs into promoters and candidate enhancers based on their transcript stability (CAGE vs PRO-cap; See methods) and the proximity to annotated (Refseq) gene transcription start sites (TSS). Based on the CAGE data, 12,878 tTREs were identified as promoters, and the remaining 74,948 tTREs were identified as enhancers (similar numbers were obtained using Refseq TSSs, see **Supplementary Table 2**). Promoters and enhancers show expected patterns of transcription initiation (**Fig. 2A**), RNA polymerase II (Pol II), H3K27 acetylation, and H3K4 methylation (**Fig. 2B**, **Supplementary Fig. 2A**). Globally, tTREs fall within accessible DNA regions and contain ENCODE annotated TFBSs, the most enriched TFBSs tending to be cell-type relevant (**2B, Supplementary Fig. 2B**) (Wang et al. 2016; Stelzer et al. 2016). The identified enhancers are enriched with regulatory information such as expression quantitative trait loci (eQTLs), chromatin accessibility QTLs (DNaseI-sensitivity QTLs, dsQTLs), and disease associated variants (GWAS SNPs) (**Fig. 2D**). Interestingly, the regulatory variants are more enriched in enhancers than promoters, which was reproduced using DNase hypersensitive sites (DHSs) to identify enhancers (**Supplementary Fig. 2C**). Together, these results demonstrate the regulatory potential of the enhancers, and the superior power of using nascent RNA sequencing to identify them.

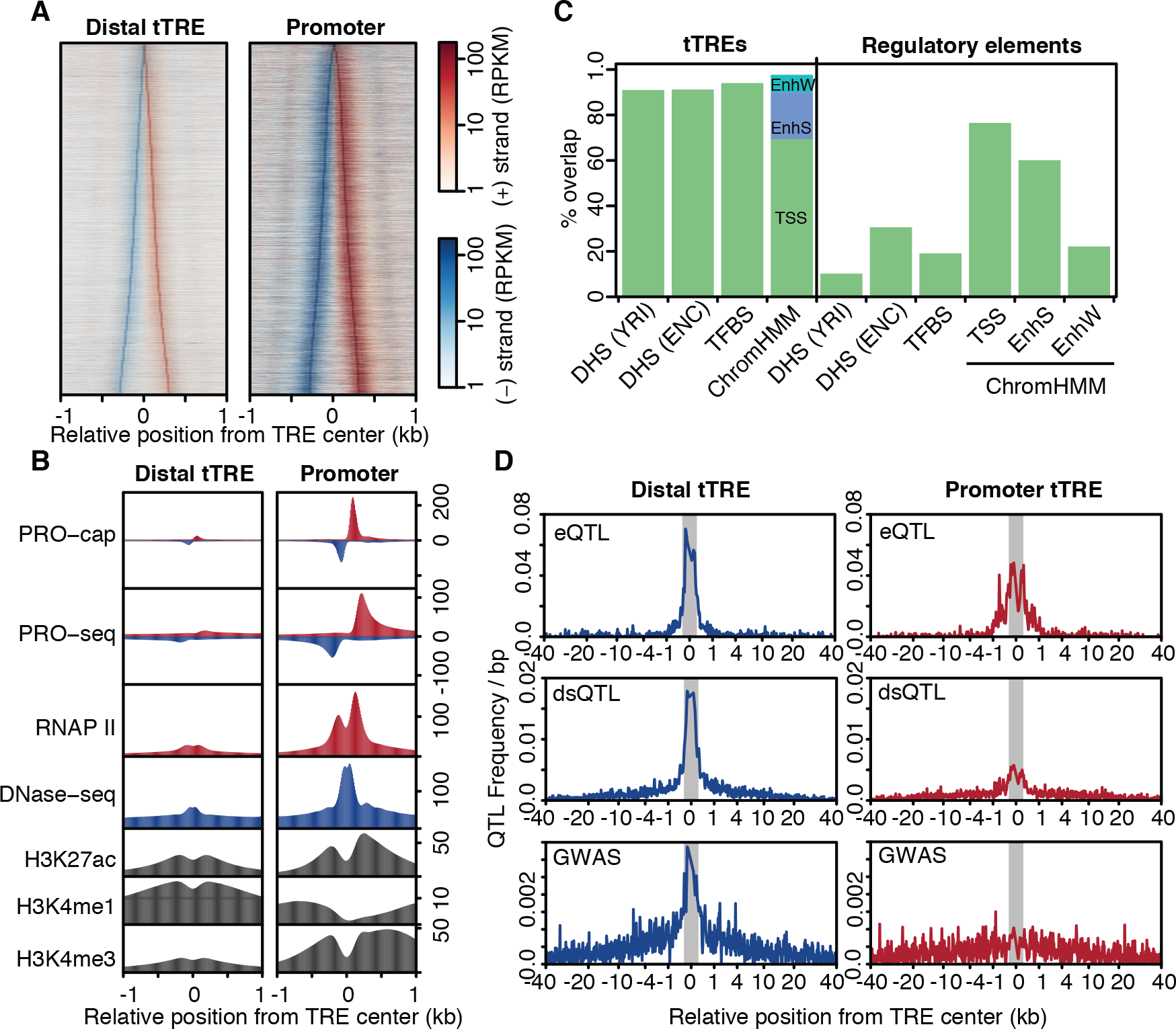
(**A**) PRO-cap signal at promoters and enhancers. Heatmap shows plus-strand (red) and minus-strand (blue) read counts at distal tTREs (putative enhancers) and promoter tTREs, ordered by increasing width. (**B**) PRO-cap-identified tTREs have characteristic promoter and enhancer chromatin patterns. Metaplots of PRO-cap, PRO-seq, DNase-seq and RNAP II, H3K27ac, H3K4me1, and H3K4me3 ChIP-seq signals at enhancers (Distal tTRE) and promoters (Promoter). Promoters are oriented in the direction of the gene. (**C)** Overlap between tTREs and regulatory regions. DHS(YRI): DNase I hypersensitive windows from the Yoruban (YRI) LCLs (n=630,168), DHS(ENC): DNase I hypersensitive sites from ENCODE LCLs (including GM12878; n=359,361), TFBS: defined by ENCODE Factorbook in GM12878 (n=212,144). ChromHMM: regions defined by ChromHMM as transcription start sites (TSS; n=21,342), strong enhancers (EnhS; n=19,362), and weak enhancers (EnhW; n=70,620) in GM12878. (**D**) PRO-cap tTREs are enriched in genetic variants associated with gene expression (eQTLs), chromatin accessibility (dsQTLs), and human disease (GWAS). Shaded regions indicate tTRE boundaries.

### Identification of genetic variants associated with transcription

To investigate how underlying sequences establish the transcriptional signature at tTREs, we tested the association between genetic variation across the individuals and the pattern of transcription at tTREs. To avoid confounding effects caused by differences in the read mappability of different alleles, we devised a variant sensitive alignment method that masks out allele-mappability biased regions (see online methods and **Supplementary Fig. 3A-B**). After allele-mappability masking, we identified 76,630 tTREs (**Supplementary Table 2**), about 40% of which are variably expressed between individuals (**Supplementary Fig. 3C-D**).

We identified genetic variants associated with the pattern of transcriptional initiation: either a change in the overall levels (overall PRO-cap read counts) or the directionality (log_2_ ratio of plus strand reads over minus strand reads, i.e. directionality index) of divergent bidirectional transcription (**Fig. 3A, B**). We mapped the genotypes to quantile-normalized PRO-cap read counts and directionality indices at tTREs within 2 kb. We named the variants associated with changes in overall PRO-cap signal transcription initiation QTLs (tiQTLs), and those associated with changes in directionality as directional initiation QTLs (diQTLs). Overall, 16,193 TREs have an associated tiQTL, and 4,162 TREs have an associated diQTL (FDR < 0.1, **Table 1**). Among those, we find that 82.7% of tiQTLs and 65.8% of diQTLs are associated with changes in transcription at enhancers rather than promoters. These numbers are comparable to QTLs affecting chromatin accessibility (dsQTL, (Degner et al. 2012)) from the same population. We validated our tiQTL results using allele specific expression analysis and estimated the average effect on tTRE transcription initiation to be around 2-fold for the most likely causal SNPs (**Supplementary Fig 4A-B**). Overall, the number and quality of QTLs allows us to observe patterns in their location within enhancers and the types of sequences they create or disrupt.

**Figure 3.**
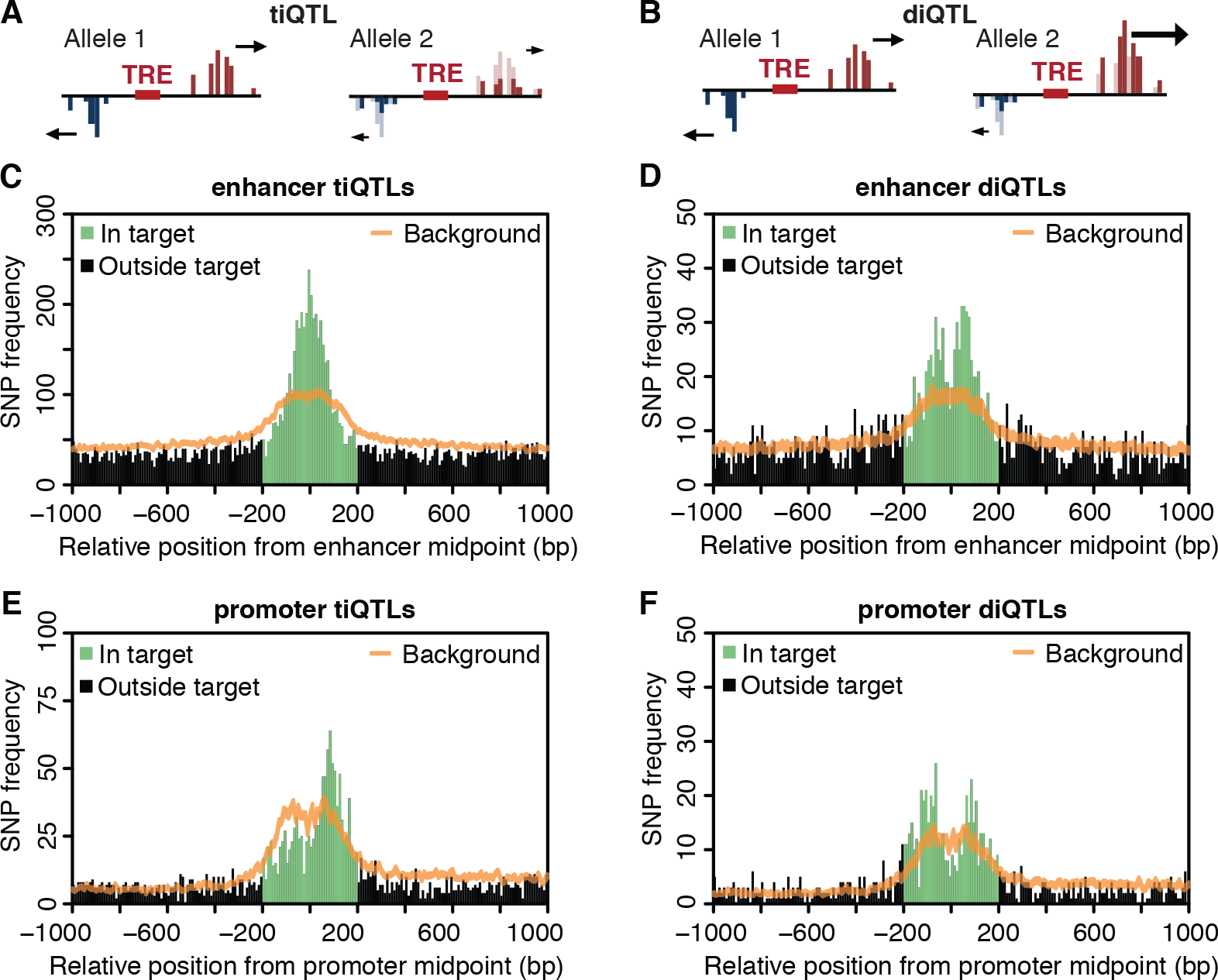
Genetic variants are associated with transcription levels and directionality at enhancers and promoters. (**A**) Transcription initiation QTL (tiQTL) schematic (**B**) Directional initiation QTL (diQTL) schematic (**C**) tiQTLs enriched at enhancer midpoints. A histogram of QTL frequency around enhancer midpoints with the expected background distribution with 99% confidence interval (sampled from all SNPs in same region) shown in orange. (**D**) diQTLs are enriched at enhancer TSSs. As in (**C**), for diQTLs. (**E**) tiQTLs are enriched at promoter TSSs. As in (**C**), at promoters except oriented so that dominant strand (usually gene) is downstream. (**F**) diQTLs are enriched at promoter TSSs. As in (**F**), for diQTLs.

We then examined how tiQTLs and diQTLs are linked to the transcriptional architecture of enhancers. To enrich for causal SNPs, we filtered ti- and diQTLs based on local minimum p-values (within 5 kb) and examined the QTL density around enhancer midpoints for both tiQTLs (**Fig. 3C**) and diQTLs (**Fig. 3D**). While both QTL types are enriched within the enhancer regions, the peak of tiQTL enrichment is at the enhancer midpoint, whereas diQTLs are most enriched ~70 bp to either side, coinciding with the average relative position of TSSs (**Supplementary Fig. 4C-D**). Based on these results, we hypothesize that the overall transcriptional activity is generally regulated from the central TF binding sites, but the directionality of transcription is mostly regulated in the core-promoter-element regions near the transcription initiation sites, where the pre-initiation complex (PIC) of RNA polymerase II assembles.

At promoters, we found that both tiQTLs and diQTLs are preferentially enriched nearer to the TSSs compared to the central region (**Fig. 3E, F**, **Supplementary Fig. 4E-F**). However, compared to enhancers, promoter-associated tiQTLs were more likely than diQTLs to be specifically enriched at the dominant strand (mostly the direction of the gene). This difference in tiQTL enrichment between enhancers and promoters suggests that the two tTRE types have different tolerances to genetic variation affecting divergent transcription.

### Transcription associated QTLs affect TF binding motifs and core promoter elements

To further explore this difference in the placement of tiQTLs and diQTLs within enhancers, we compared the underlying sequences. For example, the non-reference allele of the tiQTL rs185220, located within a proximal enhancer near the *SETD9* promoter (**Fig. 4A**), creates a perfect match to the binding site for SP1 transcription factor at the center of the associated enhancer. This alternate allele is also associated with increased eRNA transcription, which is concordant with the change in the TF binding sequence in the central region. Conversely, the non-reference allele of rs8050061 diQTL disrupts a match to the canonical Initiator element (Inr) (Ngoc et al. 2017), coinciding with decreased transcription from the affected strand (**Fig. 4B**).

**Figure 4.**
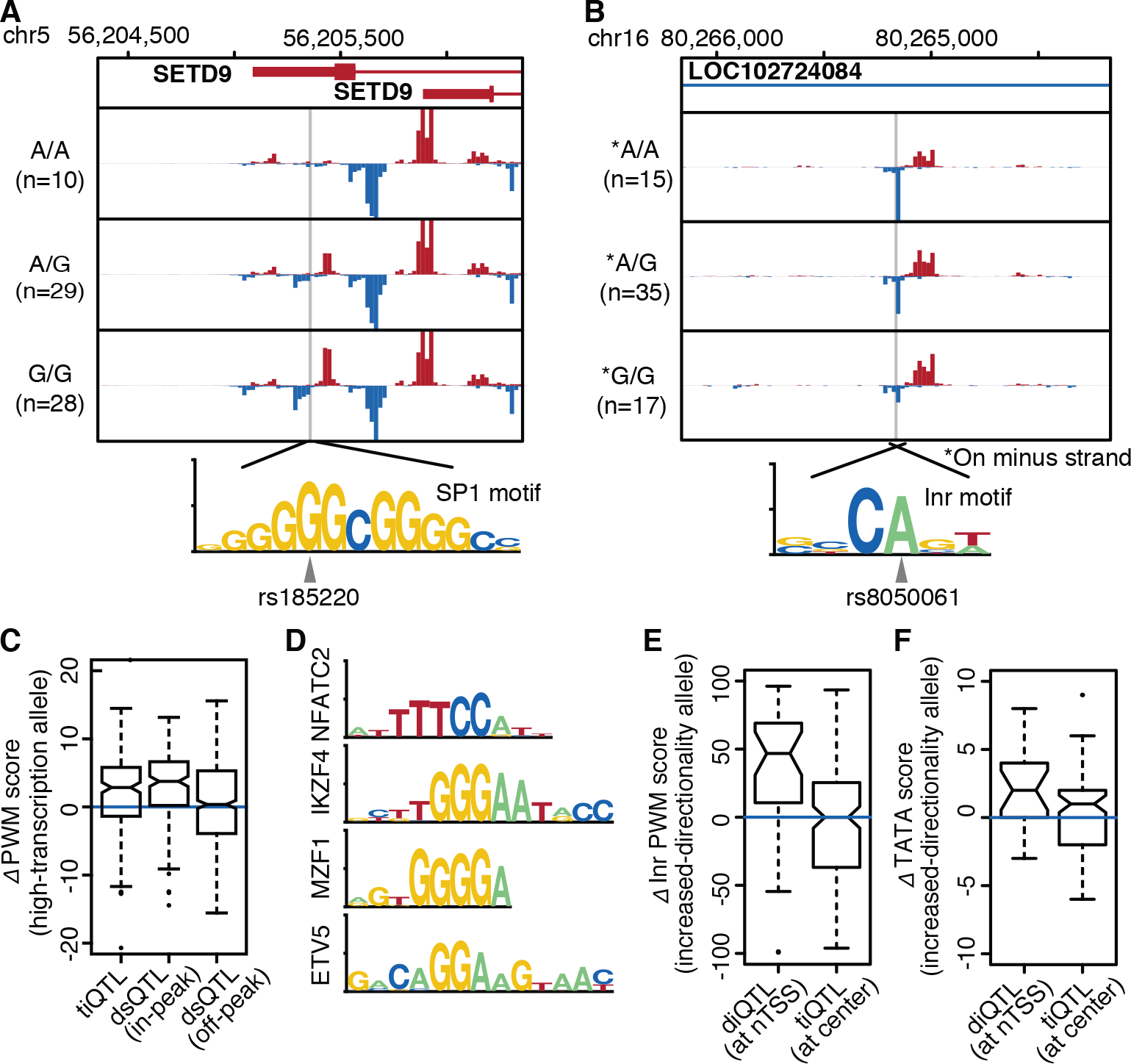
tiQTLs affect TF binding sites; diQTLs affect core promoter elements. (**A**) Average PRO-cap signal separated by genotype at tiQTL rs185220. The alternate allele creates a perfect match to the SP1 binding motif. (**B**) Average PRO-cap signal separated by genotype at diQTL rs8050061. The alternate allele disrupts a canonical human Inr motif. (**C**) Difference in PWM scores between tiQTL high transcription and low transcription alleles. Notch reflects the 95% confidence interval of the median. DNase hypersensitivity QTLs are shown for comparison. (**D**) Motifs most often affected by in-tTRE tiQTLs (>20% increase over background). (**E**) Difference in Inr match likelihood between increased and decreased directionality allele. diQTLs within 10 bp from the TSSs compared to tiQTLs within 10 bp from the tTRE center. (**F**) As in (E), for TATA-box match likelihood. Region selected for diQTLs is 40 to 20 bp upstream of TSS.

We tested the generality of the association between tiQTLs and central TF binding motifs using TFBS Position Weight Matrix (PWM) scores, as described previously (Degner et al. 2012; Weirauch et al. 2014). On average, the alleles with stronger eRNA transcription have stronger PWM score than the weaker alleles (**Fig. 4C**). The effect size is similar to what was observed in QTLs affecting chromatin accessibility (dsQTL), indicating that alteration of TF binding motifs affects both the open chromatin and eRNA transcription. The motifs most disproportionately affected are associated with immune-related TFs (**Fig. 4D**). We suspected diQTLs would affect sequence elements at regions near the enhancer TSSs that are analogous to the promoters. We defined these regions as the “core”, since they contain core promoter element-like motifs such as the Initiator element (Inr) and the TATA box. Inr is highly enriched at PRO-cap identified TSSs (Ngoc et al. 2017) (**Supplementary Fig. 5**) and diQTL alleles associated with a shift in directionality towards stronger expression of the strand where the diQTL is located have stronger matches to the motif (**Fig. 4E**). To a lesser degree, the same was also true for TATA-like elements, usually found 20-40 bp upstream of the TSS (**Fig. 4F**).

### Transcription-associated QTLs associate with gene expression

The most critical question when exploring changes in chromatin phenotypes is whether they have an effect on the final output - gene expression. We used a RNA-seq data in the same set of LCLs (Pickrell et al. 2010) to define expression QTLs (eQTLs) and test their association with tiQTLs or diQTLs. We focused on SNPs in regions immediately surrounding TREs (± 2 kb), and found 744 genes within 200 kb with associated eQTLs (FDR < 0.05). Overall, these eQTLs are enriched within 200 bp from the enhancer midpoints defined by PRO-cap (**Fig. 5A**).

We found that enhancers that contain an associated tiQTLs and/or diQTL are 8.6-fold and 13.2- fold more likely to contain an eQTL, respectively (**Supplementary Fig. 6A**). The tiQTLs or diQTLs themselves were also directly associated with gene expression (**Fig. 5B**). Importantly, diQTLs that are not tiQTLs also show this stronger association with gene expression (**Supplementary Fig. 6B**). These results indicate that both tiQTLs and diQTLs are associated with gene expression and, given the different positioning of tiQTLs and diQTLs within enhancers, that sequences at both the central TFBS and the regions surrounding the TSSs may affect enhancer function in gene regulation (**Supplementary Fig. 6C**).

**Figure 5.**
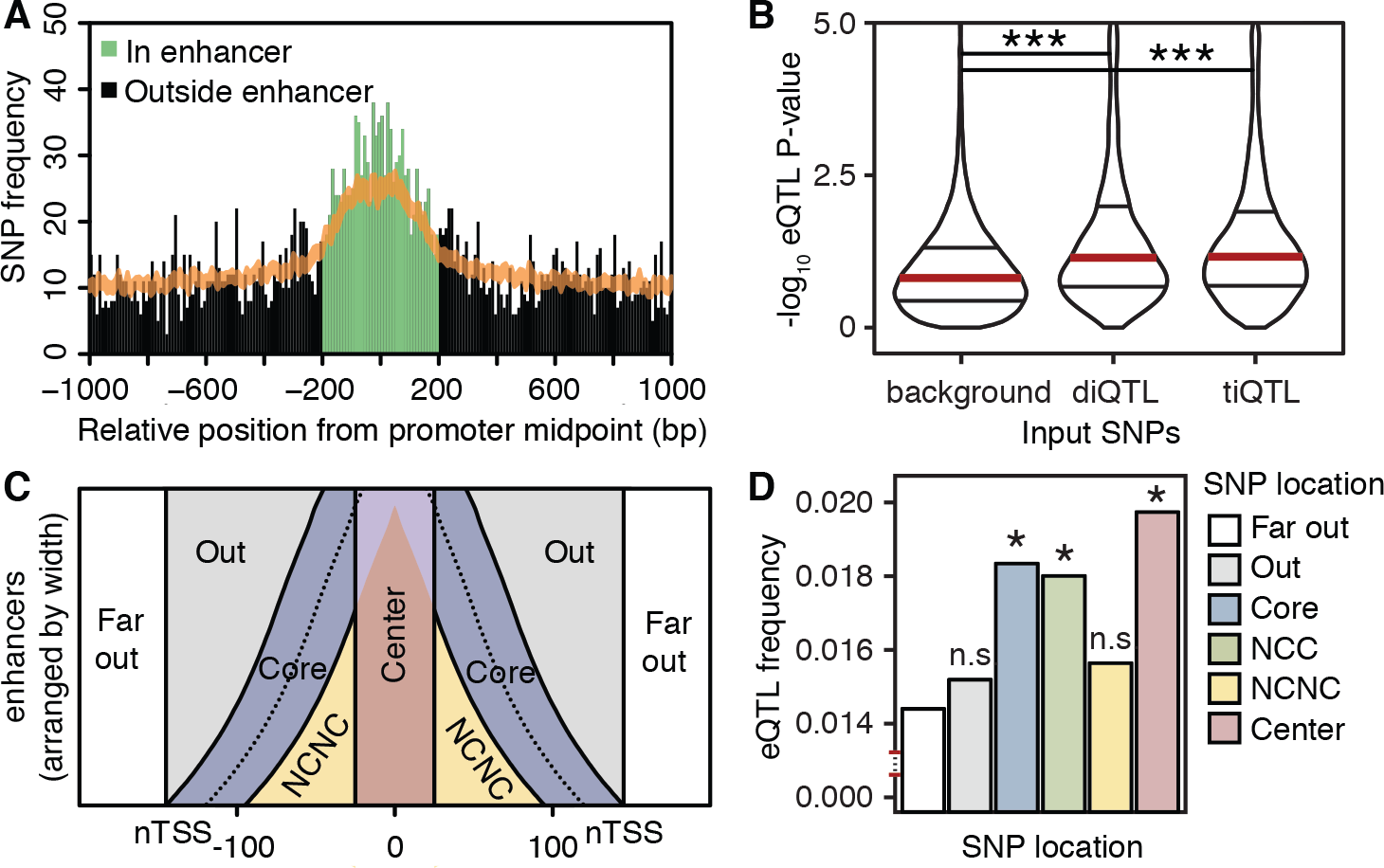
ti- and diQTLs at enhancers associate with changes in gene expression. (**A**) eQTLs are enriched at enhancers. A histogram of QTL density around enhancer midpoints. (B) Distribution of P-values for mRNA association for diQTLs, tiQTLs and background SNPs (all SNPs within 2 kb of enhancer). Violin plots are marked with quantiles, the median colored red. *** indicates P<1×10^−16^, Wilcoxon rank-sum test. (**C**) Diagram explaining the separation of candidate enhancer tTREs into functional regions. NCNC stands for non-center, non-core. (**D**) eQTL frequency in different enhancer regions. NCNC: non-center-non-core, NCC: non-center-core. Each colored bar is compared to Far Out in a Fisher’s exact test. * indicates p<0.05, n.s. indicates not-significant.

To further explore the model that both the center and TSS cores are important for enhancer function, we hypothesized that eQTLs would be enriched in those regions relative to the regions outside them. We separated enhancer regions into: the center, the core promoter-like region from which transcription arises (core), the space between them (non-core-non-center, NCNC), the space outside of the core but within the enhancer region (out), and those beyond the enhancer region (**Fig. 5C**).

Using the far-out region as a baseline, we find a significantly higher eQTL frequency within the center and the core (Fisher’s exact test, P < 0.05, **Fig. 5D**). Importantly, the core region remains high in eQTL frequency and significance when the region overlapping the center (lilac in Fig. 4C) is ignored (non-center-core: NCC). The out and NCNC regions have no significant increase in eQTL frequency. Additionally, we compared the eQTL frequency between the core and the out or NCNC regions, normalizing for the distance from the center, using bootstrapping to estimate an empirical significance level (**Supplementary Fig. 6D-F**). We find that the core regions are significantly enriched with eQTLs compared to the out regions (p < 10^-16^), and marginally enriched compared to the NCNC region (p = 0.107). These support our hypothesis that the core eRNA initiation regions are important for target gene expression, in addition to the central transcription factor binding regions.

## Discussion

We explored the activity and architecture of transcribed transcriptional regulatory elements (tTREs) by studying variation in transcription initiation across human LCLs. We identified genetic variants associated with enhancer and promoter transcriptional activity and directionality. The pattern of enrichment for these genetic variants and the types of motifs they affect suggest that overall transcriptional activity at enhancers is regulated from the central transcription-factor binding region (TFBS) and directionality is regulated from the surrounding core initiation regions. Both variant types are associated with gene expression at a higher rate than expected, indicating that both the central TFBS and the flanking core initiation regions affect transcription at enhancers and their role in gene expression. This conclusion is supported by regional enrichment of eQTLs within enhancers.

Identification of enhancers based on capped nascent RNA sequencing provides a direct measure of transcriptional activity and, therefore, shows higher sensitivity than previous methods. A direct measure of transcriptional activity is critical, as non-productive transcripts such as eRNAs are rapidly degraded in the nucleus (Andersson et al. 2014). Other transcription-based approaches, such as CAGE and nuclear short RNA analysis, are impeded by this instability. As we showed, CAGE performs well in identifying promoters but detects enhancers less efficiently than PRO-cap. Additionally, focusing on the bidirectional nascent transcription start sites (nTSSs) using PRO-cap enabled us to filter out spurious transcription from only one strand, which increases the specificity. Although it is possible that enhancer transcription can be unidirectional, our nascent transcription analysis detected mostly bidirectional and divergently paired PRO-cap peaks, and observed very few unpaired peaks.

The fact that genetic variants affecting enhancer directionality are associated with changes in gene expression brings up an interesting question of mechanism. While the change in directionality towards or away from the sense strand of the gene will obviously affect its expression levels at promoters, how directional initiation at enhancers affects gene expression is less clear. In some cases, modulating the polymerase initiation at only one of the two initiating sites could impact the overall enhancer activity, though in other cases overall transcriptional activity is not affected by the diQTLs. In the latter case, it is possible that, despite the prevailing model that enhancers are orientation-independent, there may be distance and orientation-specific effects on their target genes, potentially involving eRNAs. While our current list of diQTLs is limited due to the sample size, increasing power with a larger scale analysis would allow testing of these possibilities.

Overall, our data show the power of combining capped-nascent-RNA sequencing with human population genetics to explore the architecture of human enhancers. These results suggest a bipartite model for enhancers, where sequences at both the central TFBS and the core promoter regions surrounding the TSSs are important for enhancer function. Our model also supports the hypothesis that eRNA transcription itself can be functional. In practice, this will have an important implication for assessing the significance of disease-associated non-coding genetic variants, adding eRNA start sites as the regions of interest.

## Methods

### Preparation of PRO-cap/PRO-seq libraries from human lymphoblastoid cell lines (LCLs)

Human LCLs were acquired from the Coriell Biorepository. Unrelated individuals were selected from the Yoruban (YRI) population as described in Degner *et al.* (**Supplementary Table 1**). LCLs were cultured in RPMI 1640 media (Gibco) with 2 mM L-glutamine (Gibco) and 15% Fetal Bovine Serum (Gibco) supplements without antibiotics at 37°C under 5% CO_2_. Mid-log phase cultures were seeded at 2×10^5^ cells/ml density in 15 ml suspension culture, and maintained for 24 hr before the cell harvest. Batches of 10 randomly selected cultures were prepared at a time for a total of 100 samples (PRO-cap: 70 individuals + 10 replicates, PRO-seq: 10 individuals + 10 replicates). One individual was dropped out due to unsuccessful PRO-cap library generation and two because they lacked phased genotype data (**Supplementary Table 1**). Replicate batches, grown from independent cultures, were processed at least 2 months apart. Briefly, LCLs were pelleted by centrifugation at 800×g for 3 min in 4°C, washed twice by resuspension in 10 mL 4°C Phosphate Buffered Saline (PBS) and centrifugation at 800×g for 3 min, followed by resuspension in 50 μl of storage buffer (50 mM Tris-Cl pH 8.0, 25% glycerol, 5 mM magnesium acetate, 0.1 mM EDTA, 5 mM DTT). Cells were instantly frozen in liquid nitrogen, and stored at −80°C. Upon thawing, cells were incubated in polymerase run-on reactions with ribonucleotide triphosphate (NTP) substrates at a final concentration of 19 μM biotin-11-CTP, 19 μM biotin-11-UTP, 0.19 mM ATP, and 0.19 mM GTP at 37°C for 3 min. This was followed by nascent-RNA sequencing library preparation as described previously (Mahat et al. 2016).

### Alignment of the PRO-cap/PRO-seq reads to the reference genome

We followed a previously described PRO-seq/PRO-cap data processing procedure (Mahat et al. 2016) and combined alignments for each method (see Supplementary methods for details).

### Identification of nascent transcription start sites (nTSSs) and transcribed transcriptional regulatory elements (tTREs)

We combined all PRO-cap dataset reads and identified ~1.4 billion (1,417,065,796) unique read molecules mapped to the hg19 genome. We then used a bidirectional read count filtering approach comparable to capped RNA analysis described elsewhere to identify bidirectional transcribed tTREs (Andersson et al. 2014) (**Supplementary Figure 1A, see Supplementary Methods for details**).

### Evaluation of the sensitivity and specificity of PRO-cap

To assess the predictive power of PRO-cap to find transcriptional regulatory elements, we used the transcription start sites of the genes with greater than 1 RPKM as the TSS standards from the mRNA-seq data in the YRI LCLs (Pickrell et al. 2010) (n = 12,272), and ENCODE FACTORBOOK defined EEP300 binding sites in GM12878 cell line as the enhancer standards (n = 18,956). We used the randomly selected 1 million genomic regions in 500 bp windows as described in the previous section as a background distribution. We calculated the true positive rate as a function of different RPKM thresholds using TSS or EP300 standards, and calculated the false positive rate as the fraction of background regions above the thresholds. We generated receiver operating characteristic (ROC) curves for PRO-cap detecting TSS and EP300 sites, and compared the PRO-cap ROC with an available cap analysis of gene expression (CAGE) data in GM12878 (RIKEN CAGE; GSM849349) (**Figure 1B-C**).

### Classification of tTREs into promoters and enhancers

To classify the tTREs into gene promoters and enhancers/regulatory elements, we used two criteria: 1) distance to annotated refseq TSSs, 2) overlap with CAGE sites. For the refseq based promoter annotations, we defined the tTREs with at least one nTSSs within 500 bp of refseq TSSs as the promoters (n=14,986). We then defined tTREs greater than 2 kb away from any refseq TSSs as distal enhancers (n=34,922). For the CAGE based promoter and enhancer classifications, PRO-cap and CAGE counts at the nTSSs are collected for the plus and the minus strands separately, and RPM normalized. nTSSs with CAGE counts significantly above the background were called promoter TSSs. To estimate the background CAGE read counts, we first selected nTSSs that are at least 1 kb away from any annotated refseq TSS (n=38,658), and calculated the CAGE read counts for both the plus and the minus strands at these background regions. We calculated the p-values of the nTSS CAGE read counts based on the empirical background distribution, and found 13,833 nTSSs that have CAGE readcounts that are significantly higher than the background using FDR < 0.1 (*P* < 0.0145, CAGE RPM >= 1.18655). Of the 13,833 nTSSs, 995 were bidirectional pairs, yielding 12,878 tTREs as promoters, 995 of which are bidirectional. We defined the remaining 74,948 tTREs as enhancers.

### DNA sequence motif analysis

We used the RTFBSdb suite (Wang et al. 2016) that clusters transcription factor binding motifs based on similarity, chooses a representative motif for each cluster based on the expression data, and computes enrichment for known motifs. We filtered CIS-BP motifs based on expression in our LCLs using our PRO-seq data, clustered the motifs with agnes clustering into 400 clusters, and chose a representative motif for each cluster based on expression in our PRO-seq data. We used this motif list to look for motif enrichment within tTRE centers.

### Regulatory variant enrichment analysis

We used previously defined lists of expression QTLs and DNase I hypersensitivity QTLs and lifted the coordinates over to the hg19 genome (n=1,090 and 8,899 respectively). For the disease associated GWAS SNPs, we used the NIH GWAS Catalog entries as of March, 2015 in hg19 (n=12,239). We used the refseq annotation based distal enhancer and promoter classifications for the tTREs, and calculated the per base QTL enrichment relative to the tTRE midpoints. For comparison, we also generated enrichment plots relative to the DNase I hypsersensitivity sites (ENCODE DHS; n=52,292 promoter DHS, 226,832 distal enhancer DHS) in all LCL cell lines (**Supplementary Figure 2C**).

### Variant sensitive alignment of the PRO-cap reads to individual genomes

In summary, we reconstructed the individual phased haplotype genomes, masked out any tTRE regions with ambiguous mappability, then re-aligned the PRO-cap reads to the individual haplotype genomes (**Supplementary Figure 3A-B**). Because of removing allele mappability biased regions, we re-calculated read per million (RPM) of the read counts sum across all individuals for each tTRE, and used a RPM cut-off of 0.5 to further select tTRE peaks for testing associations (n=76,630). See supplement for details.

### Normalization of the transcription initiation phenotypes for association testing

First, we normalized the read counts to the sequencing depth. We added the plus and the minus strand read counts in each tTRE window, then divided the raw read counts by per million total read counts in the tTRE windows for each individual. Then we used a quantile normalization, where the distribution of read counts in an individual is matched to a reference distribution. For the reference distribution, we applied a “median of the ratio” normalization method that is used in the DEseq RNA-seq analysis software (Dillies et al. 2013). We used this “median of the ratio” normalized distribution for the quantile normalization of the read counts (**Supplementary Table 2**).

### Identification of variably expressed tTREs

To identify tTREs that are variably expressed, we used normalized PRO-cap read count data that contain partial replicates (n=8; **Supplementary Figure 3C**). For each tTRE, we calculated the deviation from the mean of the normalized read counts between replicates and between different samples. Then we used Wilcoxon’s rank sum test to test the alternative hypothesis that the differences between samples are greater than between the replicates for each tTRE, and calculated p-values. We estimated the number of variably expressed tTRE by analyzing the complete distribution of the p-values as described previously (Storey et al. 2007). Under the null hypothesis, p-values should have a uniform distribution with a density of 1, but the observed p-values are only uniformly distributed for large p-values. The density of the portion of the p-value distribution that is uniform is ~0.244, indicating that up to ~75.6% of tTREs can be considered variably expressed. Using an FDR of ~0.2, we identified 29,694 variably expressed nTSSs.

### Transcription initiation QTL (tiQTL) association testing

We tested the association between the nascent transcription initiation phenotypes at each tTRE region and the genotype of SNPs and short indels within a cis range of 2 kb from the midpoint of the nTSS regions. We took the variant sensitive normalized PRO-cap readcounts, and identified tiQTLs using the MatrixEQTL R package (Shabalin 2012). We used allele-mappability-bias-masked tTRE regions on autosomes (n=76,118), and variant sites with a minor allele frequency greater than 5% in our haplotype-phased individuals (n=9,808,709). A total of 994,993 pairs were tested. We tested up to 20 principal components (PCs) as co-variates in 2 kb cis-tiQTL tests and found that 16 PCs gave the largest number of significant tiQTLs (FDR>0.1). 16,193 tTREs are associated with at least one tiQTL in 2 kb cis regions.

### Directional initiation QTL (diQTL) association testing

We tested the association between the relative direction of the divergent bidirectional nascent transcription initiation pairs at the two nTSSs in each tTRE, and the genotypes of the genetics variants. To compute a metric for the directionality of tTREs, we calculated the directionality index as ratio of plus-strand (1 - 250) and minus-strand (−250 - 0) read counts, log_2_ transformed the ratio and quantile normalized the resulting index using the same method we used for diQTLs. The association of these directionality phenotypes with genotype was assessed, as for tiQTLs, using MatrixEQTL. We determined that using 8 principal components as covariates in association testing gave the largest number of significant associations.

### Local-minimum p-value filtering of QTLs for more likely causal SNPs

We split the genome into 5 kb windows, staggered by 1 kb so that each SNP is in 5 different 5 kb windows. In each window we keep only the SNP with the lowest *P*-value, independent of which tTRE or gene it affects. If there were two or more SNPs with the lowest P-value, none were kept.

### Measuring the effect of tiQTL SNPs on TF sequence motifs

We tested the effect of tiQTLs on TF binding likelihood as described by Degner et al. (*13*). We limited our analysis to the tiQTLs at the center-region (midpoint ±40 bp) of tTREs and imposed a stricter filtering requirement by keeping only those QTLs where the most significantly associated SNP has a P-value an order of magnitude lower than the next most significant. Reference and alternative alleles were categorized into “enhancing” and “repressive” alleles based on the relative PRO-cap readcounts around the tTREs. We used CIS-BP human transcription factor frequency matrices to generate position weight matrices (PWMs), and queried a 40 bp region surrounding each tiQTL for strong matches to the PWMs (motif score > 13). We repeated this analysis for both alleles and compared the resulting motif scores for enhancing and repressive alleles. We then took these motif score differences in these strong-matched TF motifs to obtain the overall effect of each variation on TF binding potential. For comparison, we also performed this analysis on dsQTLs (Degner et al. 2012) that fall within the most highly affected window (100 bp windows) and dsQTLs outside the affected windows.

### Identifying motif enrichment among motifs disrupted by tiQTLs

We used motifbreakR (Coetzee et al. 2015) to calculate the motif disruption score for each tiQTL for the 400 motifs selected from the CIS-BP motif database with RTFBSdb. A motif is considered disrupted by the tiQTL if there is a significant match to a motif in at least one allele (p < 0.01) and there is at least a 0.5 bit difference in motif matching scores between the alleles. We then count the number of times each motif is disrupted by a tiQTL and compare it to the number of times that same motif is disrupted by randomly selected SNPs within the same region (+/- 200 bp from tTRE midpoint).

### Measuring the effect of diQTL SNPs on Initiator elements

Initiator (Inr) element likelihood is calculated using published human Inr frequency matrix (Ngoc et al. 2017) by taking the natural exponent of the PWM scores. First, we selected diQTL SNPs within 5 base pairs (bp) from the defined nascent transcription start sites (nTSSs) of tTREs. We calculated the strand specific Inr likelihood difference between the two alleles and included diQTLs that generate differences in Inr likelihood score (>5). We oriented the diQTL directionality effect sizes towards the strand in which the Inr scores are calculated, and assigned the Inr likelihood difference as (high directional allele - low directionality allele). For comparison, we selected tiQTL SNPs within 5 bp from the center of tTREs, calculated Inr likelihood differences in between relatively higher directional allele to the lower directional allele towards the direction of the Inr element, and plotted the distribution of ΔInr likelihood similar to diQTLs.

### Measuring the effect of diQTL SNPs on TATA-like elements

We focused on the area 40 to 20 bp upstream of tTRE TSSs and looked for instances of TATA, or inversions thereof. We oriented the diQTLs as explained for Inr and calculated a delta TATA score (high directionality - low directionality allele). A perfect TATA got a score of 4, a single inversion (A -> T, T -> A) such as TTTA or TAAA got a score of 2, and two inversions got a score of 1. Anything else scored 0.

### Expression QTL association testing

Using only SNPs that fall within 2 kb of TREs (same range as the ti- and diQTLs), we tested the association between SNPs and gene expression. By limiting our analysis only to the region we are interested in, we increase our power to detect significant associations. We used RNA-seq data from Pickrell et al. (*11*), merged replicates by taking an average across replicates for each gene, and performed the same quantile normalization as above. We used matrixEQTL to identify eQTLs within a cis-distance of 200 kb (fdr < 0.05). We determined that using 13 PCs as covariates gave the largest number of significant associations.

### Overlap between eQTL- and diQTL/tiQTL-containing tTREs

We identified enhancers that contain associated diQTLs and/or tiQTLs within 200 bps of the enhancer midpoint. We then computed the proportion of those that overlap an eQTL and compared that ratio with the proportion of enhancers without such QTLs that overlap an eQTL.

### Gene expression association of tiQTLs/diQTLs

We computed a p-value for the association of each filtered tiQTL and diQTL with gene expression using matrixEQTL and a cis-distance of 200 kb, reporting all p-values. We then compared the distribution of p-values (-log10) with those for all SNPs within 2 kb of TREs.

### Regional eQTL enrichment analysis

Enhancers were split into regions according to figure 4C. Center is +/- 25 bp from the midpoint and core is +/- 25 bp from the TSS. The number of eQTLs in each region was counted and normalized to the total number of SNPs in the region.

### Data access

The sequencing libraries from this study have been submitted to the NCBI Gene Expression Omnibus (GEO) with accession number GSE110638. Custom scripts available upon request.

## Acknowledgements

We thank Dr. Jun Hee Lee from the University of Michigan, Dr. John Lis and Dr. Charles Danko from Cornell University, and members of their labs, for helpful discussions during the inception and completion of this work. This work was supported a CU-CVG scholars award (to KK).

## Author contributions

Project was conceived of by H.K. and H.M.K. Cell culture, library preparations, sequencing, and processing of raw data were done by H.K. in H.M.K.’s lab. K.K., H.K., N.D.T., and H.M.K. performed the analyses and generated figures. K.K., H.K., Y.K., H.M.K., and J.T.L. wrote the manuscript.

## References

Andersson R, Gebhard C, Miguel-Escalada I, Hoof I, Bornholdt J, Boyd M, Chen Y, Zhao X, Schmidl C, Suzuki T, et al. 2014. An atlas of active enhancers across human cell types and tissues. Nature 507: 455–461.

Andersson R, Sandelin A, Danko CG. 2015. A unified architecture of transcriptional regulatory elements. Trends Genet 31: 426–433.

Banovich NE, Lan X, McVicker G, Geijn B van de, Degner JF, Blischak JD, Roux J, Pritchard JK, Gilad Y. 2014. Methylation QTLs Are Associated with Coordinated Changes in Transcription Factor Binding, Histone Modifications, and Gene Expression Levels. PLOS Genetics 10: e1004663.

Battle A, Khan Z, Wang SH, Mitrano A, Ford MJ, Pritchard JK, Gilad Y. 2015. Genomic variation. Impact of regulatory variation from RNA to protein. Science 347: 664–667.

Cannavò E, Koelling N, Harnett D, Garfield D, Casale FP, Ciglar L, Gustafson HE, Viales RR, Marco-Ferreres R, Degner JF, et al. 2017. Genetic variants regulating expression levels and isoform diversity during embryogenesis. Nature 541: 402–406.

Coetzee SG, Coetzee GA, Hazelett DJ. 2015. motifbreakR: an R/Bioconductor package for predicting variant effects at transcription factor binding sites. Bioinformatics 31: 3847–3849.

Core LJ, Martins AL, Danko CG, Waters CT, Siepel A, Lis JT. 2014. Analysis of nascent RNA identifies a unified architecture of initiation regions at mammalian promoters and enhancers. Nat Genet 46: 1311–1320.

Degner JF, Pai AA, Pique-Regi R, Veyrieras J-B, Gaffney DJ, Pickrell JK, De Leon S, Michelini K, Lewellen N, Crawford GE, et al. 2012. DNase I sensitivity QTLs are a major determinant of human expression variation. Nature 482: 390–394.

Dillies M-A, Rau A, Aubert J, Hennequet-Antier C, Jeanmougin M, Servant N, Keime C, Marot G, Castel D, Estelle J, et al. 2013. A comprehensive evaluation of normalization methods for Illumina high-throughput RNA sequencing data analysis. Brief Bioinformatics 14: 671–683.

ENCODE Project Consortium. 2012. An integrated encyclopedia of DNA elements in the human genome. Nature 489: 57–74.

Ferreira PG, Oti M, Barann M, Wieland T, Ezquina S, Friedländer MR, Rivas MA, Esteve-Codina A, GEUVADIS Consortium, Rosenstiel P, et al. 2016. Sequence variation between 462 human individuals fine-tunes functional sites of RNA processing. Sci Rep 6: 32406.

Garieri M, Delaneau O, Santoni F, Fish RJ, Mull D, Carninci P, Dermitzakis ET, Antonarakis SE, Fort A. 2017. The effect of genetic variation on promoter usage and enhancer activity. Nat Commun 8: 1358.

Gusev A, Lee SH, Trynka G, Finucane H, Vilhjálmsson BJ, Xu H, Zang C, Ripke S, Bulik-Sullivan B, Stahl E, et al. 2014. Partitioning heritability of regulatory and cell-type-specific variants across 11 common diseases. Am J Hum Genet 95: 535–552.

Hah N, Murakami S, Nagari A, Danko CG, Kraus WL. 2013. Enhancer transcripts mark active estrogen receptor binding sites. Genome Res 23: 1210–1223.

Henriques T, Scruggs BS, Inouye MO, Muse GW, Williams LH, Burkholder AB, Lavender CA, Fargo DC, Adelman K. 2018. Widespread transcriptional pausing and elongation control at enhancers. Genes Dev 32: 26–41.

Kaikkonen MU, Spann NJ, Heinz S, Romanoski CE, Allison KA, Stender JD, Chun HB, Tough DF, Prinjha RK, Benner C, et al. 2013. Remodeling of the Enhancer Landscape during Macrophage Activation Is Coupled to Enhancer Transcription. Molecular Cell 51: 310–325.

Kasowski M, Grubert F, Heffelfinger C, Hariharan M, Asabere A, Waszak SM, Habegger L, Rozowsky J, Shi M, Urban AE, et al. 2010. Variation in Transcription Factor Binding Among Humans. Science 328: 232–235.

Kim T-K, Hemberg M, Gray JM, Costa AM, Bear DM, Wu J, Harmin DA, Laptewicz M, Barbara-Haley K, Kuersten S, et al. 2010. Widespread transcription at neuronal activity-regulated enhancers. Nature 465: 182–187.

Kruesi WS, Core LJ, Waters CT, Lis JT, Meyer BJ. 2013. Condensin controls recruitment of RNA polymerase II to achieve nematode X-chromosome dosage compensation. Elife 2: e00808.

Kwak H, Fuda NJ, Core LJ, Lis JT. 2013. Precise maps of RNA polymerase reveal how promoters direct initiation and pausing. Science 339: 950–953.

Lam MTY, Li W, Rosenfeld MG, Glass CK. 2014. Enhancer RNAs and regulated transcriptional programs. Trends Biochem Sci 39: 170–182.

Li YI, van de Geijn B, Raj A, Knowles DA, Petti AA, Golan D, Gilad Y, Pritchard JK. 2016. RNA splicing is a primary link between genetic variation and disease. Science 352: 600–604.

Mahat DB, Kwak H, Booth GT, Jonkers IH, Danko CG, Patel RK, Waters CT, Munson K, Core LJ, Lis JT. 2016. Base-pair-resolution genome-wide mapping of active RNA polymerases using precision nuclear run-on (PRO-seq). Nat Protoc 11: 1455–1476.

Majewski J, Pastinen T. 2011. The study of eQTL variations by RNA-seq: from SNPs to phenotypes. Trends Genet 27: 72–79.

Maurano MT, Humbert R, Rynes E, Thurman RE, Haugen E, Wang H, Reynolds AP, Sandstrom R, Qu H, Brody J, et al. 2012. Systematic localization of common disease-associated variation in regulatory DNA. Science 337: 1190–1195.

McVicker G, van de Geijn B, Degner JF, Cain CE, Banovich NE, Raj A, Lewellen N, Myrthil M, Gilad Y, Pritchard JK. 2013. Identification of genetic variants that affect histone modifications in human cells. Science 342: 747–749.

Ngoc LV, Cassidy CJ, Huang CY, Duttke SHC, Kadonaga JT. 2017. The human initiator is a distinct and abundant element that is precisely positioned in focused core promoters. Genes Dev 31: 6–11.

Pickrell JK, Marioni JC, Pai AA, Degner JF, Engelhardt BE, Nkadori E, Veyrieras J-B, Stephens M, Gilad Y, Pritchard JK. 2010. Understanding mechanisms underlying human gene expression variation with RNA sequencing. Nature 464: 768–772.

Schor IE, Degner JF, Harnett D, Cannavò E, Casale FP, Shim H, Garfield DA, Birney E, Stephens M, Stegle O, et al. 2017. Promoter shape varies across populations and affects promoter evolution and expression noise. Nature Genetics 49: ng.3791.

Shabalin AA. 2012. Matrix eQTL: ultra fast eQTL analysis via large matrix operations. Bioinformatics 28: 1353–1358.

Stelzer G, Rosen N, Plaschkes I, Zimmerman S, Twik M, Fishilevich S, Stein TI, Nudel R, Lieder I, Mazor Y, et al. 2016. The GeneCards Suite: From Gene Data Mining to Disease Genome Sequence Analyses. Curr Protoc Bioinformatics 54: 1.30.1–1.30.33.

Storey JD, Madeoy J, Strout JL, Wurfel M, Ronald J, Akey JM. 2007. Gene-expression variation within and among human populations. Am J Hum Genet 80: 502–509.

Wang Z, Martins AL, Danko CG. 2016. RTFBSDB: an integrated framework for transcription factor binding site analysis. Bioinformatics 32: 3024–3026.

Weirauch MT, Yang A, Albu M, Cote AG, Montenegro-Montero A, Drewe P, Najafabadi HS, Lambert SA, Mann I, Cook K, et al. 2014. Determination and inference of eukaryotic transcription factor sequence specificity. Cell 158: 1431–1443.

